# Addressing Overlapping Sample Challenges in Genome-Wide Association Studies: Meta-Reductive Approach

**DOI:** 10.1101/2023.12.08.570867

**Authors:** Farid Rajabli

## Abstract

Polygenic risk scores (PRS) are instrumental in genetics, offering insights into an individual level genetic risk to a range of diseases based on accumulated genetic variations. These scores rely on Genome-Wide Association Studies (GWAS). However, precision in PRS is often challenged by the requirement of extensive sample sizes and the potential for overlapping datasets that can inflate PRS calculations. In this study, we present a novel methodology, Meta-Reductive Approach (MRA), that was derived algebraically to adjust GWAS results, aiming to neutralize the influence of select cohorts. Our approach recalibrates summary statistics using algebraic derivations. Validating our technique with datasets from Alzheimer’s disease studies, we showed perfect correlation between summary statistics of proposed approach and “leave-one-out” strategy. This innovative method offers a promising avenue for enhancing the accuracy of PRS, especially when derived from meta-analyzed GWAS data.

**Availability and implementation:** MRA script is freely available at https://github.com/hihg-um/MRA as an R function.

## Introduction

Polygenic risk scores (PRS) have emerged as an essential tool in the field of genetics ^1,2^. These scores offer a unique insight into an individual’s genetic predisposition to a wide array of diseases and traits, capturing the cumulative effects of multiple genetic variants^3^. The Genome-Wide Association Studies (GWAS) serve as the base for creating PRS ^4^. GWAS investigates the entire genetic makeup of individuals to identify genetic variations associated with specific diseases or traits. The predictive accuracy and precision of PRS are enhanced when the base GWAS summary statistics come from a sizeable sample, and the population in the GWAS matches the population where the PRS is being applied ^4,5^. Due to this need for a substantial sample size, studies often aim to meta-analyze all available genetic datasets to achieve the statistical power necessary for identifying genetic markers linked to the trait or disease. However, this approach presents a challenge in securing independent datasets for training, testing, and validating PRS performance ^6^. The use of overlapping samples can inflate the PRS calculations, resulting in imprecise risk predictions.

A logical approach might be to exclude a specific cohort of interest and then rerun meta-analyses with the remaining datasets. However, given the significant computational resources needed and the difficulties in accessing detailed summary statistics for all cohorts, this isn’t always viable. Nonetheless, we do have access to the cohort-level data for the specific dataset we aim to employ as a training and testing set. Recognizing this advantage, we formulated an alternative technique that incorporates the cohort-level result of our chosen dataset along with the meta-analysis GWAS findings. The goal is to neutralize the impact of the overlapping cohort of interest on the meta-analysis GWAS summary statistics, thus producing a PRS that avoids the inflationary tendencies arising from overlapping samples.

In this study, we derived equations to adjust GWAS results, effectively eliminating the impact of selected cohorts. Through comprehensive simulations and real data analysis, we demonstrated that our methodology effectively updates the base data’s summary statistics, thereby addressing the challenge.

### Derivation of Adjusted Summary Statistics: Meta-Reductive Approach

We analyzed two distinct sets of summary statistics:

1. A compilation from *n* datasets meta-analyzed using an inverse variance-based approach ^7^.
2. A specific dataset of interest that was also part of the meta-analysis.

For these datasets:

- B and *SE* symbolize the effect size and standard error, respectively, from the aggregate meta-analysis across *n* datasets.
- *β* _*i*_ and *se*_*i*_ specify the effect size and standard error for the individual cohort *i*.

Our primary aim was to compute a summary statistic that eliminates the influence of the dataset of interest, providing a clearer perspective on the overarching genetic structure.

### i. Inverse-Variance-Weighted Effect-Size Estimation

The inverse variance method gives more weight to studies with smaller variance because they offer more precise estimates. The weight, *w*_*i*_, is the inverse of the variance, or squared standard error, of the effect size, β_*i*_.

Given,

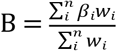 where the 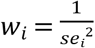

Expanding this:

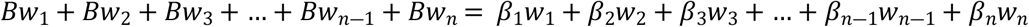

This is the weighted sum of the effect sizes across all datasets, including the one of interest. Now, to remove the effect of the specific dataset, β_*n*_, we rearrange:

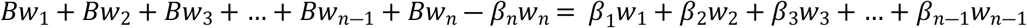

Which yields:

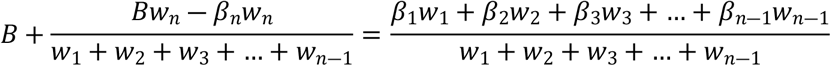

This equation essentially adjusts the overall effect size, *B*, by subtracting the influence of the dataset of interest.

### ii. Standard Error Derivation

The standard error (SE) offers a measure of the statistical accuracy of an estimate. Here, we adjust the SE based on the weights of all datasets excluding the one of interest.

Using:

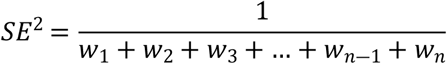

We derive:

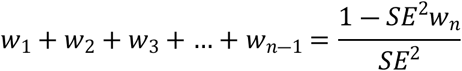

This equation gives the combined weight of all datasets, excluding the dataset of interest.

### iii. Adjusted Effect Size and Standard Error

Post removing the influence of the dataset of interest, the modified effect size is given by:

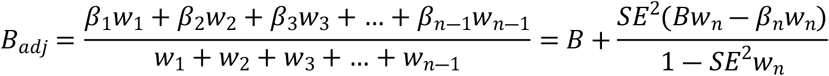

This adjusted beta, *B*_*adj*_, having nullified the contribution of the specific dataset *n*. Additionally, the adjusted standard error is:

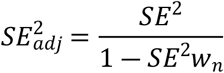

This adjustment ensures that the standard error reflects the precision of our new effect size estimate, free from the influence of the specific dataset.

### Validation

To validate our methodological approach, we utilized summary statistics from four publicly accessible Alzheimer disease studies: Kunkle et al.^8^, Kunkle et al.^9^ AA, Bellinguez et al.^10^, and Moreno-Grau S. et al. ^11^ From these studies, 100,000 markers were selected to conduct a meta-analysis using the METASOFT software^12^.

Following the initial meta-analysis, we applied a systematic “leave-one-out” strategy. For each iteration, we excluded the summary statistics from one dataset and conducted a meta-analysis of the remaining three. The results from this procedure served as our individual-level data for the three datasets in question.

For the final step of validation, we calculated the adjusted *B*_*adj*_ and 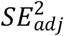 values based on our proposed method and compared them against the individual-level data derived from the “leave-one-out” meta-analyses. Our results showed perfect positive correlation between summary statistics of three datasets using “leave-one-out” strategy and our approach with adjusted *B*_*adj*_ and 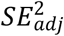 values. Figure 1 illustrates this correlation for both effect size and standard error.

**Figure 1.**
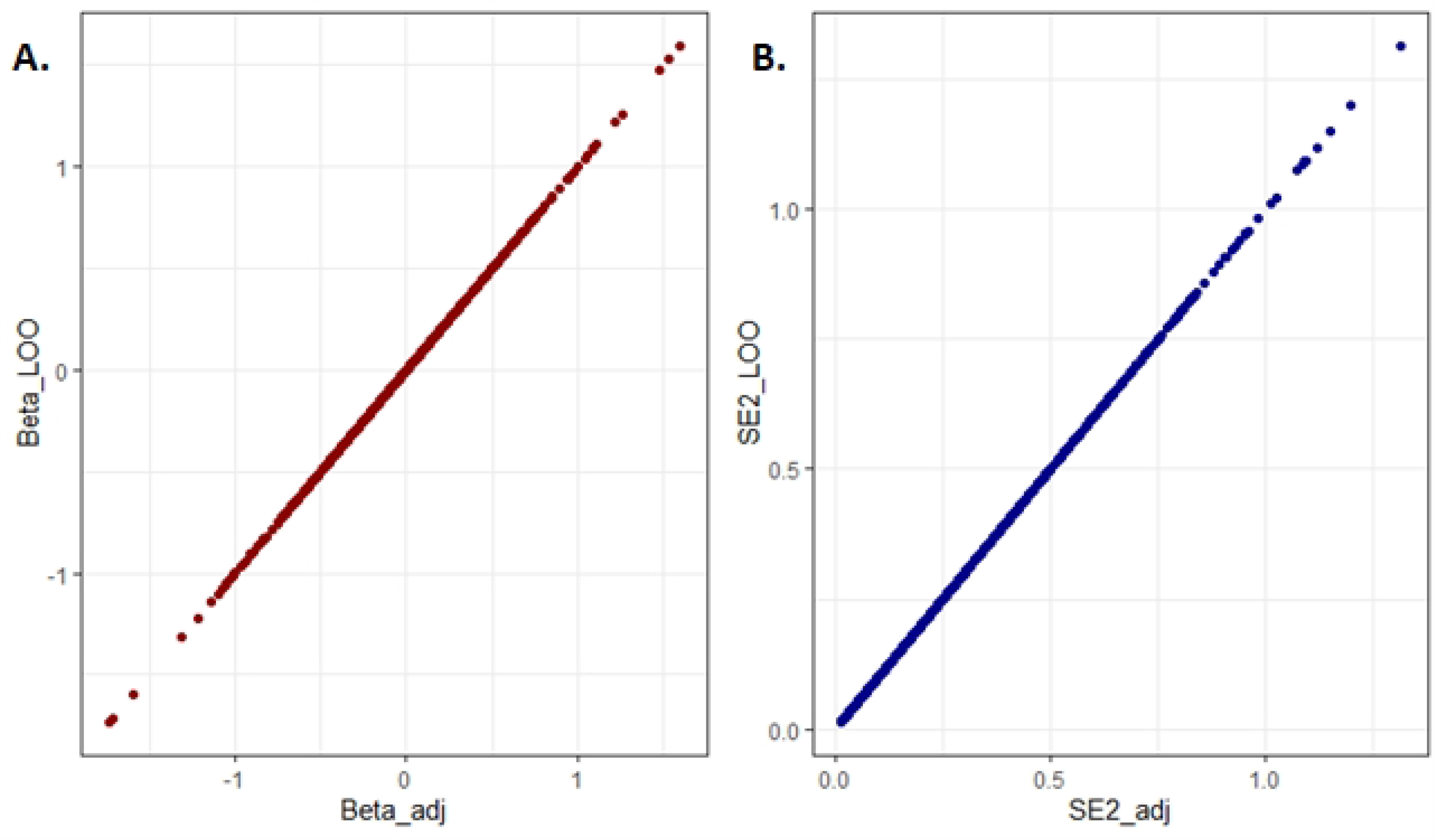
Comparison between the adjusted results from the Meta-Reductive Analysis (MRA) (Beta_adj and SE2_adj) approach and the “leave one out” (Beta_LOO and SE2_LOO). The MRA-adjusted values show perfect positive correlation with the “leave one out” calculation for both Beta values (A) and Standard Error (B).

## Ethical Approval

No ethical statement was needed, as in this study, publicly available summary statistics were used.

## Author Contributions

Farid Rajabli contributed to conceptualization, methodology, software, validation, formal analysis, investigation, data curation, writing—original draft preparation, and visualization.

## References

1. Padilla-Martinez, F., Collin, F., Kwasniewski, M., and Kretowski, A. (2020). Systematic Review of Polygenic Risk Scores for Type 1 and Type 2 Diabetes. Int. J. Mol. Sci. 5, 10.3390/ijms21051703.

2. Harrison, J.R., Mistry, S., Muskett, N., and Escott-Price, V. (2020). From Polygenic Scores to Precision Medicine in Alzheimer’s Disease: A Systematic Review. Journal of Alzheimer’s disease 4, 1271–1283.

3. Gallagher, S., Hughes, E., Wagner, S., Tshiaba, P., Rosenthal, E., Roa, B.B., Kurian, A.W., Domchek, S.M., Garber, J., Lancaster, J., Weitzel, J.N., Gutin, A., Lanchbury, J.S., and Robson, M. (2020). Association of a Polygenic Risk Score With Breast Cancer Among Women Carriers of High- and Moderate-Risk Breast Cancer Genes. JAMA Netw. Open 7, e208501.

4. Choi, S.W., Mak, T.S., and O’Reilly, P.F. (2020). Tutorial: a guide to performing polygenic risk score analyses. Nat. Protoc. 9, 2759–2772.

5. Anonymous Genome-wide association studies | Nature Reviews Methods Primers.

6. Choi, S.W., Mak, T.S.H., Hoggart, C.J., and O’Reilly, P.F. (2022). EraSOR: a software tool to eliminate inflation caused by sample overlap in polygenic score analyses. Gigascience.

7. Willer, C.J., Li, Y., and Abecasis, G.R. (2010). METAL: fast and efficient meta-analysis of genomewide association scans. Bioinformatics 17, 2190–2191.

8. Kunkle, B.W., Grenier-Boley, B., Sims, R., Bis, J.C., Damotte, V., Naj, A.C., van der Lee, S., Ahmad, S., Adams, H., Vojinovic, D., Comic, H., Roshchupkin, G., Rivadeneira, F., Uitterlinden, A., Amin, N., Ikram, A., Duijn, C., Lambert, J., and Pericak-Vance, M.A. (2019). Genetic meta-analysis of diagnosed Alzheimer’s disease identifies new risk loci and implicates A beta, tau, immunity and lipid processing. Nature genetics 3, 414-+.

9. Kunkle, B.W., Schmidt, M., Klein, H.U., Naj, A.C., Hamilton-Nelson, K.L., Larson, E.B., Evans, D.A., De Jager, P.L., Crane, P.K., Buxbaum, J.D., Ertekin-Taner, N., Barnes, L.L., Fallin, M.D., Manly, J.J., Go, R.C.P., Obisesan, T.O., Kamboh, M.I., Bennett, D.A., Hall, K.S., Goate, A.M., Foroud, T.M., Martin, E.R., Wang, L.S., Byrd, G.S., Farrer, L.A., Haines, J.L., Schellenberg, G.D., Mayeux, R., Pericak-Vance, M.A., Reitz, C., Alzheimer’s Disease Genetics Consortium, (., Graff-Radford, N.R., Martinez, I., Ayodele, T., Logue, M.W., Cantwell, L.B., Jean-Francois, M., Kuzma, A.B., Adams, L.D., Vance, J.M., Cuccaro, M.L., Chung, J., Mez, J., Lunetta, K.L., Jun, G.R., Lopez, O.L., Hendrie, H.C., Reiman, E.M., Kowall, N.W., Leverenz, J.B., Small, S.A., Levey, A.I., Golde, T.E., Saykin, A.J., Starks, T.D., Albert, M.S., Hyman, B.T., Petersen, R.C., Sano, M., Wisniewski, T., Vassar, R., Kaye, J.A., Henderson, V.W., DeCarli, C., LaFerla, F.M., Brewer, J.B., Miller, B.L., Swerdlow, R.H., Van Eldik, L.J., Paulson, H.L., Trojanowski, J.Q., Chui, H.C., Rosenberg, R.N., Craft, S., Grabowski, T.J., Asthana, S., Morris, J.C., Strittmatter, S.M., and Kukull, W.A. (2021). Novel Alzheimer Disease Risk Loci and Pathways in African American Individuals Using the African Genome Resources Panel: A Meta-analysis. JAMA Neurol. 1, 102–113.

10. Bellenguez, C., Kucukali, F., Jansen, I.E., Kleineidam, L., Moreno-Grau, S., Amin, N., Naj, A.C., Campos-Martin, R., Grenier-Boley, B., Andrade, V., Holmans, P.A., Boland, A., Damotte, V., van der Lee, S.J., Costa, M.R., Kuulasmaa, T., Yang, Q., de Rojas, I., Bis, J.C., Yaqub, A., Prokic, I., Chapuis, J., Ahmad, S., Giedraitis, V., Aarsland, D., Garcia-Gonzalez, P., Abdelnour, C., Alarcon-Martin, E., Alcolea, D., Alegret, M., Alvarez, I., Alvarez, V., Armstrong, N.J., Tsolaki, A., Antunez, C., Appollonio, I., Arcaro, M., Archetti, S., Pastor, A.A., Arosio, B., Athanasiu, L., Bailly, H., Banaj, N., Baquero, M., Barral, S., Beiser, A., Pastor, A.B., Below, J.E., Benchek, P., Benussi, L., Berr, C., Besse, C., Bessi, V., Binetti, G., Bizarro, A., Blesa, R., Boada, M., Boerwinkle, E., Borroni, B., Boschi, S., Bossu, P., Brathen, G., Bressler, J., Bresner, C., Brodaty, H., Brookes, K.J., Brusco, L.I., Buiza-Rueda, D., Burger, K., Burholt, V., Bush, W.S., Calero, M., Cantwell, L.B., Chene, G., Chung, J., Cuccaro, M.L., Carracedo, A., Cecchetti, R., Cervera-Carles, L., Charbonnier, C., Chen, H.H., Chillotti, C., Ciccone, S., Claassen, J.A.H.R., Clark, C., Conti, E., Corma-Gomez, A., Costantini, E., Custodero, C., Daian, D., Dalmasso, M.C., Daniele, A., Dardiotis, E., Dartigues, J.F., de Deyn, P.P., de Paiva Lopes, K., de Witte, L.D., Debette, S., Deckert, J., Del Ser, T., Denning, N., DeStefano, A., Dichgans, M., Diehl-Schmid, J., Diez-Fairen, M., Rossi, P.D., Djurovic, S., Duron, E., Duzel, E., Dufouil, C., Eiriksdottir, G., Engelborghs, S., Escott-Price, V., Espinosa, A., Ewers, M., Faber, K.M., Fabrizio, T., Nielsen, S.F., Fardo, D.W., Farotti, L., Fenoglio, C., Fernandez-Fuertes, M., Ferrari, R., Ferreira, C.B., Ferri, E., Fin, B., Fischer, P., Fladby, T., Fliessbach, K., Fongang, B., Fornage, M., Fortea, J., Foroud, T.M., Fostinelli, S., Fox, N.C., Franco-Macias, E., Bullido, M.J., Frank-Garcia, A., Froelich, L., Fulton-Howard, B., Galimberti, D., Garcia-Alberca, J.M., Garcia-Gonzalez, P., Garcia-Madrona, S., Garcia-Ribas, G., Ghidoni, R., Giegling, I., Giorgio, G., Goate, A.M., Goldhardt, O., Gomez-Fonseca, D., Gonzalez-Perez, A., Graff, C., Grande, G., Green, E., Grimmer, T., Grunblatt, E., Grunin, M., Gudnason, V., Guetta-Baranes, T., Haapasalo, A., Hadjigeorgiou, G., Haines, J.L., Hamilton-Nelson, K.L., Hampel, H., Hanon, O., Hardy, J., Hartmann, A.M., Hausner, L., Harwood, J., Heilmann-Heimbach, S., Helisalmi, S., Heneka, M.T., Hernandez, I., Herrmann, M.J., Hoffmann, P., Holmes, C., Holstege, H., Vilas, R.H., Hulsman, M., Humphrey, J., Biessels, G.J., Jian, X., Johansson, C., Jun, G.R., Kastumata, Y., Kauwe, J., Kehoe, P.G., Kilander, L., Stahlbom, A.K., Kivipelto, M., Koivisto, A., Kornhuber, J., Kosmidis, M.H., Kukull, W.A., Kuksa, P.P., Kunkle, B.W., Kuzma, A.B., Lage, C., Laukka, E.J., Launer, L., Lauria, A., Lee, C.Y., Lehtisalo, J., Lerch, O., Lleo, A., Longstreth, W., Jr, Lopez, O., de Munain, A.L., Love, S., Lowemark, M., Luckcuck, L., Lunetta, K.L., Ma, Y., Macias, J., MacLeod, C.A., Maier, W., Mangialasche, F., Spallazzi, M., Marquie, M., Marshall, R., Martin, E.R., Montes, A.M., Rodriguez, C.M., Masullo, C., Mayeux, R., Mead, S., Mecocci, P., Medina, M., Meggy, A., Mehrabian, S., Mendoza, S., Menendez-Gonzalez, M., Mir, P., Moebus, S., Mol, M., Molina-Porcel, L., Montrreal, L., Morelli, L., Moreno, F., Morgan, K., Mosley, T., Nothen, M.M., Muchnik, C., Mukherjee, S., Nacmias, B., Ngandu, T., Nicolas, G., Nordestgaard, B.G., Olaso, R., Orellana, A., Orsini, M., Ortega, G., Padovani, A., Paolo, C., Papenberg, G., Parnetti, L., Pasquier, F., Pastor, P., Peloso, G., Perez-Cordon, A., Perez-Tur, J., Pericard, P., Peters, O., Pijnenburg, Y.A.L., Pineda, J.A., Pinol-Ripoll, G., Pisanu, C., Polak, T., Popp, J., Posthuma, D., Priller, J., Puerta, R., Quenez, O., Quintela, I., Thomassen, J.Q., Rabano, A., Rainero, I., Rajabli, F., Ramakers, I., Real, L.M., Reinders, M.J.T., Reitz, C., Reyes-Dumeyer, D., Ridge, P., Riedel-Heller, S., Riederer, P., Roberto, N., Rodriguez-Rodriguez, E., Rongve, A., Allende, I.R., Rosende-Roca, M., Royo, J.L., Rubino, E., Rujescu, D., Saez, M.E., Sakka, P., Saltvedt, I., Sanabria, A., Sanchez-Arjona, M.B., Sanchez-Garcia, F., Juan, P.S., Sanchez-Valle, R., Sando, S.B., Sarnowski, C., Satizabal, C.L., Scamosci, M., Scarmeas, N., Scarpini, E., Scheltens, P., Scherbaum, N., Scherer, M., Schmid, M., Schneider, A., Schott, J.M., Selbaek, G., Seripa, D., Serrano, M., Sha, J., Shadrin, A.A., Skrobot, O., Slifer, S., Snijders, G.J.L., Soininen, H., Solfrizzi, V., Solomon, A., Song, Y., Sorbi, S., Sotolongo-Grau, O., Spalletta, G., Spottke, A., Squassina, A., Stordal, E., Tartan, J.P., Tarraga, L., Tesi, N., Thalamuthu, A., Thomas, T., Tosto, G., Traykov, L., Tremolizzo, L., Tybjaerg-Hansen, A., Uitterlinden, A., Ullgren, A., Ulstein, I., Valero, S., Valladares, O., Broeckhoven, C.V., Vance, J., Vardarajan, B.N., van der Lugt, A., Dongen, J.V., van Rooij, J., van Swieten, J., Vandenberghe, R., Verhey, F., Vidal, J.S., Vogelgsang, J., Vyhnalek, M., Wagner, M., Wallon, D., Wang, L.S., Wang, R., Weinhold, L., Wiltfang, J., Windle, G., Woods, B., Yannakoulia, M., Zare, H., Zhao, Y., Zhang, X., Zhu, C., Zulaica, M., EADB, GR@ACE, DEGESCO, EADI, GERAD, Demgene, FinnGen, ADGC, CHARGE, Farrer, L.A., Psaty, B.M., Ghanbari, M., Raj, T., Sachdev, P., Mather, K., Jessen, F., Ikram, M.A., de Mendonca, A., Hort, J., Tsolaki, M., Pericak-Vance, M.A., Amouyel, P., Williams, J., Frikke-Schmidt, R., Clarimon, J., Deleuze, J.F., Rossi, G., Seshadri, S., Andreassen, O.A., Ingelsson, M., Hiltunen, M., Sleegers, K., Schellenberg, G.D., van Duijn, C.M., Sims, R., van der Flier, W.M., Ruiz, A., Ramirez, A., and Lambert, J.C. (2022). New insights into the genetic etiology of Alzheimer’s disease and related dementias. Nat. Genet. 4, 412–436.

11. Moreno-Grau, S., de Rojas, I., Hernandez, I., Quintela, I., Montrreal, L., Alegret, M., Hernandez-Olasagarre, B., Madrid, L., Gonzalez-Perez, A., Maronas, O., Rosende-Roca, M., Mauleon, A., Vargas, L., Lafuente, A., Abdelnour, C., Rodriguez-Gomez, O., Gil, S., Santos-Santos, M.A., Espinosa, A., Ortega, G., Sanabria, A., Perez-Cordon, A., Canabate, P., Moreno, M., Preckler, S., Ruiz, S., Aguilera, N., Pineda, J.A., Macias, J., Alarcon-Martin, E., Sotolongo-Grau, O., GR@ACE consortium, DEGESCO consortiums, Alzheimer’s Disease Neuroimaging Initiative, Marquie, M., Monte-Rubio, G., Valero, S., Benaque, A., Clarimon, J., Bullido, M.J., Garcia-Ribas, G., Pastor, P., Sanchez-Juan, P., Alvarez, V., Pinol-Ripoll, G., Garcia-Alberca, J.M., Royo, J.L., Franco, E., Mir, P., Calero, M., Medina, M., Rabano, A., Avila, J., Antunez, C., Real, L.M., Orellana, A., Carracedo, A., Saez, M.E., Tarraga, L., Boada, M., and Ruiz, A. (2019). Genome-wide association analysis of dementia and its clinical endophenotypes reveal novel loci associated with Alzheimer’s disease and three causality networks: The GR@ACE project. Alzheimers Dement. 10, 1333–1347.

12. Han, B., and Eskin, E. (2011). Random-Effects Model Aimed at Discovering Associations in Meta-Analysis of Genome-wide Association Studies. American journal of human genetics 5, 586–598.

